# Neural signatures of engagement and event segmentation during story listening in background noise

**DOI:** 10.1101/2025.07.15.664921

**Authors:** Björn Herrmann, Aysha Motala, Ryan A. Panela, Ingrid S. Johnsrude

## Abstract

Speech in everyday life is often masked by background noise, making comprehension effortful. Characterizing brain activity patterns when individuals listen to masked speech can help clarify the mechanisms underlying such effort. However, most previous research has focused on neural activity related to short, disconnected sentences that little resemble the more continuous, story-like spoken speech individuals typically encounter. In the current study, we used functional magnetic resonance imaging (fMRI) in humans (both sexes) to investigate how neural signatures of story listening change in the presence of masking by 12-talker babble noise. We show that, as speech masking increases, spatial and temporal activation patterns in auditory regions become more idiosyncratic to each listener. In contrast, spatial (and to some extent temporal) activity patterns in brain networks linked to effort (e.g. cinguloopercular network involving the anterior insula and anterior cingulate) are more similar across listeners when speech is highly masked and less intelligible, suggesting shared neural processes. Moreover, at times during stories when one meaningful event ended and another began, neural activation increased over extensive regions in frontal, parietal, and medial cortices. This event-boundary response appeared little affected by background noise, suggesting that listeners process meaningful units and, in turn, the gist in naturalistic, continuous speech even when it is masked somewhat by background noise. Overall, the current data may indicate that people stay engaged and cognitive processes associated with naturalistic speech processing remain intact under moderate levels of background noise, whereas auditory processing becomes more idiosyncratic to each listener.

**Significance statement:** This functional imaging study investigates how neural signatures of story listening change in the presence of masking multi-talker babble noise. We show that, as speech masking increases, auditory activity patterns become more idiosyncratic to each listener. Spatial activation patterns in effort-related regions (anterior insula, cingulate) become more similar across listeners when speech is strongly masked, indicating shared neural representations. Neural activation also increased at times when one meaningful story event ended and another began. This event-boundary response appeared little affected by moderate levels of background noise, suggesting that listeners process meaningful units in continuous speech even as background babble reduces speech intelligibility. The data indicate that many cognitive processes associated with listening to spoken stories remain intact under background noise.

## Introduction

Speech in everyday life is often degraded or masked by background noise, which makes comprehension cognitively demanding and effortful (Eckert et al., 2016; Pichora-Fuller et al., 2016; Peelle, 2018; Herrmann and Johnsrude, 2020b). Listening effort is considered an early sign of age-related hearing loss (Pichora-Fuller et al., 2016; Helfer and Jesse, 2021) and characterizing what happens in the brain when individuals listen to degraded or masked speech may clarify the mechanisms underlying listening effort (Eckert et al., 2016; Johnsrude and Rodd, 2016). Research thus far has focused mainly on the neural processes related to speech degradation/masking while individuals listen to short, disconnected sentences (Obleser and Kotz, 2010; Okada et al., 2010; Wild et al., 2012b; Ritz et al., 2022) that little resemble the more continuous, story-like spoken speech individuals often encounter (Jefferson, 1978; Mullen and Yi, 1995; Bohanek et al., 2009). Continuous speech requires a listener to mentally organize the speech stream into meaningful units that span across sentences and enables shared immersive experiences such as suspense, anticipation, empathy, and enjoyment (Speer et al., 2004; Whitney et al., 2009; Michelmann et al., 2021). Such critical speech processes are not captured when individual sentences – that do not follow a coherent narrative – are used as stimuli.

Research using functional magnetic resonance imaging (fMRI) to study the responses to sentences masked with noise has shown activations in the cingulo-opercular network (e.g., cingulate cortex, insula), pre-frontal cortex, and parietal networks as degradation/masking of speech increases and listening becomes more effortful, whereas activity in the anterior and posterior temporal cortex increases as speech becomes more intelligible (Davis and Johnsrude, 2003; Scott et al., 2006; Obleser and Kotz, 2010; Wild et al., 2012b; Scott and McGettigan, 2013; Evans et al., 2016; Ritz et al., 2022). Whether the activation patterns observed for degraded/masked sentences also hold for more naturalistic, continuous speech listening is unclear.

Other fMRI work has focused on neural processes during the perception of continuous, naturalistic stimuli, such as movies and spoken, clear speech (Nummenmaa et al., 2014; Chen et al., 2017; Regev et al., 2019; Hamilton and Huth, 2020). When materials are engaging, neural activity in wide networks, including the default mode and fronto-parietal networks, synchronizes across observers or listeners (Honey et al., 2012; Nummenmaa et al., 2014; Chen et al., 2017; Nguyen et al., 2019; Regev et al., 2019). This neural synchronization, indexed as inter-subject correlation (ISC) of neural activity (Hasson et al., 2010; Nastase et al., 2019), increases as immersive engagement with and shared understanding of the materials increase (Schmälzle et al., 2015; Nguyen et al., 2019; Song et al., 2021). Behavioral data and scalp-recorded electroencephalography (EEG) further suggest that engagement with spoken stories may be unaffected by moderate background noise (Herrmann and Johnsrude, 2020a; Irsik et al., 2022a; Yasmin et al., 2023). However, scalp EEG is ill-suited to disentangle activity from different neural systems that may differ in the nature and magnitude of shared activity and how these change when speech comprehension is effortful due to background noise. Here, we aim to identify regions in which activity becomes more idiosyncratic (less synchronized) across listeners as speech masking increases, reflecting individual responses to increasing listening challenges. We also aim to identify regions that show increased synchronization with speech masking, reflecting a common response.

Although natural environments unfold continuously, individuals mentally organize them into discrete, temporally extended events that reflect the gist of information over several tens of seconds to minutes (Zacks et al., 2007; Zacks and Swallow, 2007). Individuals tend to agree on when one event ends and another one begins, henceforth referred to as an event boundary (Kurby and Zacks, 2008; Richmond et al., 2017; Sasmita and Swallow, 2022; Michelmann et al., 2025), and this agreement is consistent with the shared neural activity patterns across individuals observed through ISC (Hasson et al., 2004; Hasson et al., 2010; Nastase et al., 2019). The accurate mental organization of natural environments into meaningful events is associated with better recall of relevant information at a later time (Zacks et al., 2006; Sargent et al., 2013; Kurby and Zacks, 2018; Newberry and Bailey, 2019). Critically, neural activity transiently increases around the time of an event boundary, potentially reflecting the increased processing demands associated with updating mental representations at an event boundary (Speer et al., 2007; Whitney et al., 2009; Zacks et al., 2010; Ben-Yakov and Henson, 2018; Barnett et al., 2024). How neural activity associated with event-boundary processing changes when individuals listen to spoken stories under varying degrees of background masking noise is unknown, but if it reflects overarching story comprehension then it should be somewhat robust to speech masking (Yasmin et al., 2023).

In the current fMRI study, we analyze blood-oxygenation level-dependent (BOLD) signal to investigate how neural signatures of engagement and event segmentation change while individuals listen to naturalistic, spoken stories masked by different degrees of background babble that degrades intelligibility but may not affect comprehension as much. Analyses focus on masking-related changes in overall activation, inter-subject synchronization of spatial and temporal patterns of brain activity, and neural responses at the times of event boundaries.

## Methods and materials

### Participants

Forty adults participated in the current study (median age: 23 years, age range: 17–34 years; 16 male, 24 female). Data from 5 additional individuals were recorded but excluded from data analysis because a few volumes during functional imaging were not recorded (n=1) or behavioral performance was at chance level, suggesting inattentive listening to the spoken stories (n=4). Participants were native English speakers or learned English before the age of 5 years. Participants reported having no hearing impairment. Participants gave written informed consent prior to the experiment and were paid $20 CAD per half-hour for their participation. The study was conducted in accordance with the Declaration of Helsinki, the Canadian Tri-Council Policy Statement on Ethical Conduct for Research Involving Humans (TCPS2-2014), and was approved by the Research Ethics Board of the University of Western Ontario.

### Acoustic stimuli and procedures

The experiment was run using PsychToolBox (version 3.0.14) on a Lenovo ThinkPad W550s laptop under Windows 7. Visual stimuli were presented to participants in the MR scanner through a mirror system. Acoustic stimuli were presented via a Steinberg UR22 external sound card and played to participants through Sensimetrics MR compatible headphones (model S14). Auditory stimuli were presented at a comfortable listening level, determined at the beginning of the fMRI session by playing a ∼1 min story in the scanner.

Participants listened to three stories from the story-telling podcast ‘The Moth’ (themoth.org). The selected stories were *“Reach for the stars one small step at a time”* by Richard Garriott (∼13 min), *“The bounds of comedy”* by Colm O’Regan (∼10 min), and *“Nacho challenge”* by Omar Qureshi (∼11 min). The Moth stories are about human experiences and life events, and they are intended to create an engaging and enjoyable listening experience. The Moth stories mirror speech in everyday life, such as disfluencies, filler-words, sentence fragments, corrections, unintentional pauses, and more flexible grammar (Tree, 1995; Bortfeld et al., 2001; Panela et al., 2024). The Moth stories are increasingly used in behavioral and neuroimaging studies due to their naturalness (Ki et al., 2016; Simony et al., 2016; Regev et al., 2019; Herrmann and Johnsrude, 2020a; Irsik et al., 2022a, b; Panela et al., 2024).

Twelve-talker babble from the Revised Speech in Noise test (Bilger, 1984; Bilger et al., 1984) was added to the stories at different speech-clarity conditions. Specifically, the speech-clarity level changed pseudo-randomly every 30-33 s among five conditions: clear speech, +14 dB, +9 dB, +4 dB, - 1 dB signal-to-noise ratio (SNR; Figure 1), such that a particular speech-clarity condition could not repeat immediately. In a previous study using the same stories, speech intelligibility ranged from 95 to 55 percent correctly heard words for clear speech and speech at +12 dB, +7 dB, +2 dB, -3 dB SNR (Irsik et al., 2022a). To achieve somewhat similar intelligibility levels in the current study, we increased the SNR levels by 2 dB relative to the previous work because the MRI scanner generates minor additional background noise. SNR was manipulated by adjusting the sound level of both the story and the babble masker to ensure that the overall sound level remained constant throughout a story and across stories. Four 30-s and 33-s segments per speech-clarity condition were presented for stories by Colm O’Regan and Omar Qureshi, respectively, and five 32-s segments per speech-clarity condition were presented for the story by Richard Garriott.

**Figure 1:**
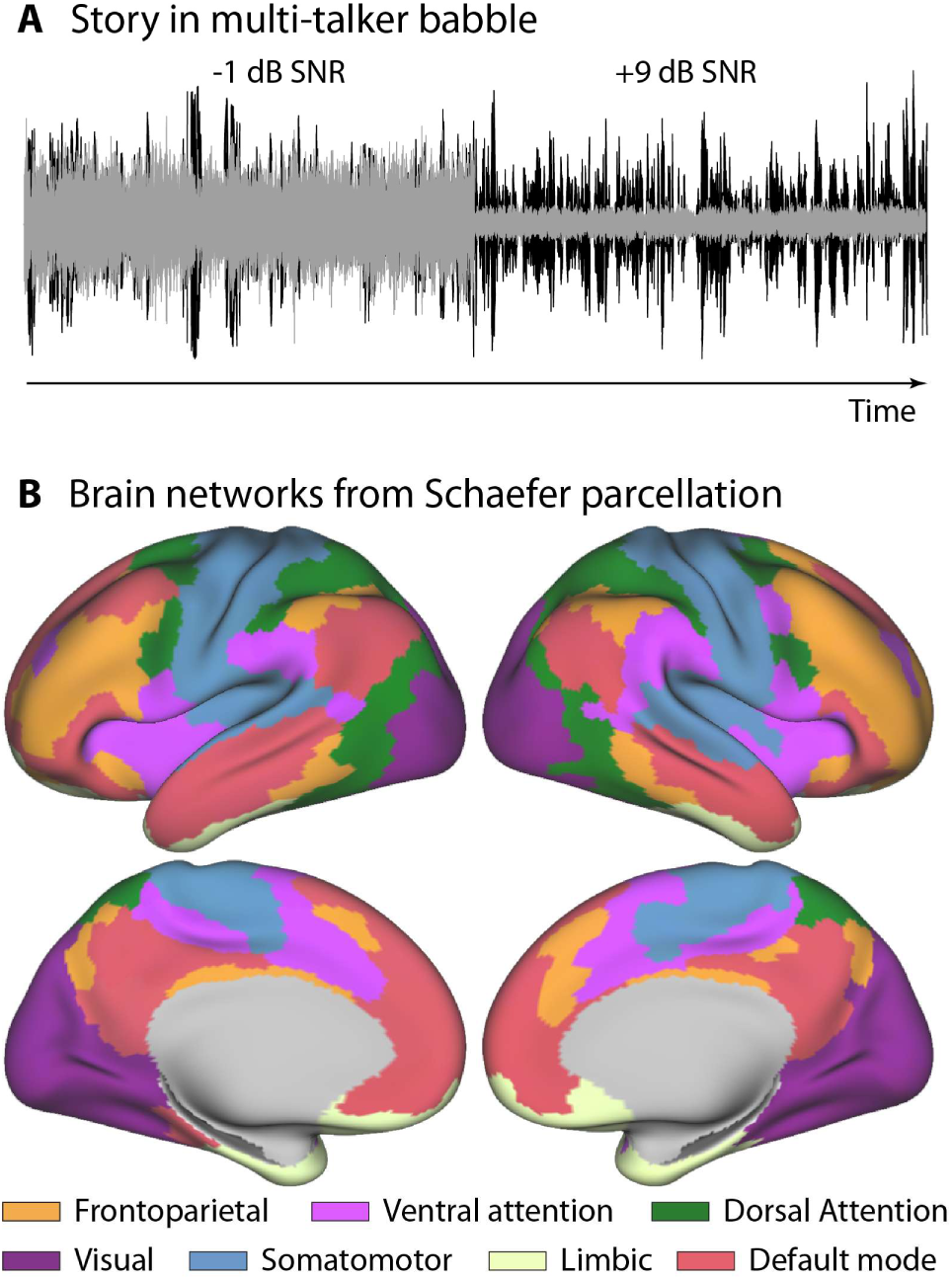
Illustration of the story-in-noise stimulation and brain parcellation. **A:** A story was played continuously and the signal-to-noise ratio of the speech to the babble masker changed every 30-33 s seconds. For better visualization, the speech and babble are displayed separately (but were added for the experiment). Speech is displayed in black. The babble masker is displayed in gray. **B:** Brain parcellation and grouping into 7 networks of the Schaefer atlas (Yeo et al., 2011; Schaefer et al., 2018).

The current study aimed to investigate inter-subject correlation of neural activity (Hasson et al., 2010; Nastase et al., 2019), which requires that each participant listens to the same story segment masked by the same speech-clarity level (Regev et al., 2019; Irsik et al., 2022a). The randomization of speech-clarity levels was thus fixed across participants for each story. To ensure that specific parts of a story were not confounded with a specific speech-clarity condition, three different versions of speech-clarity randomization were created for each story, and participants were assigned to one of the three versions. The order in which stories were presented was counterbalanced across participants.

After each story, participants performed a brief story-comprehension task to assess whether participants had attentively listened. Eight statements were visually presented, and participants had to indicate via button press whether the statement was correct or incorrect. The proportion of correct responses was calculated and data from a participant were excluded if their proportion of correct responses was at or below chance level (0.5) for any of the stories. Data from four participants were excluded for this reason.

### Recording of MRI data

Magnetic resonance imaging (MRI) was conducted using a 3T Siemens MAGNETOM Prisma Fit scanner (Siemens, Erlangen, Germany) with a 32-channel head-coil at the Centre for Functional and Metabolic Mapping at the University of Western Ontario. Participants were comfortably positioned in the bore and wore air-conduction headphones (Sensimetrics S14).

Functional images, using multi-band echo-planar imaging (Feinberg et al., 2010; Moeller et al., 2010; Barth et al., 2016) with an acceleration factor of 3, were acquired in 48 slices (interleaved), covering most of the brain, including temporal, frontal, parietal, and occipital cortices (subcortical regions and the cerebellum were not included systematically). The sequence was set up with an echo time (TE) of 30 ms, a flip angle of 40 degrees, a repetition time (TR) of 1.0 s. The matrix was 84 × 84 pixels (7/8 partial phase) with a field of view of 208 mm^2^. The in-plane resolution was 2.5 × 2.5 mm^2^. The slice thickness was 2.5 mm. Functional images were acquired in three independent runs which were separated by a short break of about 30–60 s in which scanning was discontinued. The number of volumes acquired were 608, 676, and 811 for the story by Colm O’Regan, Omar Qureshi, and Richard Garriott, respectively. The number of volumes per story included ∼5 s (5 volumes) after the story ended to account for the slow temporal evolution of the hemodynamic response function during data analysis (Buckner, 1998; Lindquist et al., 2009; Taylor et al., 2018).

For each participant, a T_1_-weighted anatomical image of the brain was recorded for co-registration during data pre-processing. The T_1_-weighted image was acquired using the following parameters: TR 2.3 s, TE = 2.98 ms, TI = 0.9 s, number of slices = 176, matrix = 256 × 256, field of view = 256 mm^2^, and a voxel size of 1 mm × 1 mm × 1 mm.

### Preprocessing of data

Data were preprocessed using SPM12 (Friston, 2007; Welcome Trust Centre for Neuroimaging, London, UK) and custom MATLAB scripts. For each of the 3 functional runs per participant, the preprocessing comprised slice-time correction (accounting for multi-band slice acquisition), rigid-body spatial realignment and unwarping, segmentation of the T_1_-weighted image, co-registration to the T_1_-weighted image according to spatial normalization parameters from segmentation, normalization to the Montreal Neurological Institute (MNI) space (MNI152NLin2009cAsym; Fonov et al., 2009), and interpolation to a 2 mm × 2 mm × 2 mm voxel size. Spatial smoothing was not applied because most analyses focused on the averaged BOLD signal within anatomically defined brain regions (essentially smoothing across voxels). For those analyses that focused on the voxel-wise BOLD signal, we were interested in the spatial activation patterns across voxels within a brain region and thus wanted to avoid reducing potentially relevant variability. Motion-related artifacts were addressed by regressing out the six standard motion parameter timeseries, which include translation (x, y, z) and rotation (roll, pitch, yaw), using the 3dDetrend function from AFNI (Analysis of Functional NeuroImages) software (Cox, 1996; Cox and Hyde, 1997). Preprocessing resulted in three 4D functional datasets, one for each story, that were used for subsequent data analysis.

### Data reduction to regions of whole-brain parcellation

To reduce data dimensionality and computational times, all data analyses focused on regions from a whole-brain parcellation. In the current study, the Schaefer atlas was used to obtain 500 brain regions per hemisphere (Schaefer et al., 2018), which are further grouped into 7 networks (Figure 1B; Yeo et al., 2011; Schaefer et al., 2018). The Schaefer atlas was chosen because the regions are of approximately similar size, making this atlas particularly useful for dimensionality reduction while covering the whole cortex. For most analyses, BOLD activity time courses from different voxels within a region were averaged to obtain one time course per Schaefer region. The averaged time course for each region was the basis for subsequent analyses (temporal inter-subject correction; event-boundary analysis). The only exception was the analysis of spatial inter-subject correlation, for which the BOLD signal was averaged across time separately for each voxel, and analyses focused on the mean BOLD signal for each voxel within a Schaefer region (see below).

To visualize BOLD signal time courses for some analyses (see event-boundary analyses below), response time courses were further averaged across individual Schaefer regions, separately for each of the 7 networks (Figure 1B; Yeo et al., 2011; Schaefer et al., 2018). The details for each specific analysis are described in the next section.

### Data and statistical analysis

#### BOLD activity to the amplitude of the speech envelope

Our initial data analysis focused on BOLD responses in each region of the Schaefer atlas to the acoustic properties of the speech signal (Honey et al., 2012; Rowland et al., 2018). The amplitude envelope was obtained for each story by calculating the absolute value of the clear speech signal. To match the sampling frequency of the BOLD signal (1 s TR), the mean amplitude envelope within 1-s time bins was calculated for each story. The resampled amplitude envelope was convolved with a canonical hemodynamic response function (Lindquist et al., 2009; Honey et al., 2012), separately for each story. A design matrix was created that contained a unique regressor for each story’s convolved envelope, a unique intercept regressor for each story, and an overall intercept regressor. One general linear model was calculated (Friston et al., 1994) for each participant and region of the Schaefer atlas, using the BOLD activity time course (concatenated for the three stories) as the predicted variable and the design matrix as the predictor. A contrast vector was multiplied with the estimated coefficients, such that the story regressors were set to 1 and the other regressors to 0. This resulted in one contrast coefficient for each participant and region of the Schaefer atlas. Positive values indicate a larger BOLD signal with increasing speech amplitude, whereas negative values indicate a larger BOLD signal with decreasing speech amplitude. For each Schaefer region, contrast coefficients were tested against zero using a one-sample t-test (group-level analysis) and the resulting t-value was converted to a z-score. Z-scores were mapped onto a partially inflated standard brain surface and visualized using the workbench environment of the Human Connectome Project (Marcus et al., 2011). Z-scores were thresholded at 3.89, corresponding to a Bonferroni-corrected significance threshold (alpha value of 0.05 divided by 1000 regions, converted to a z-score equals 3.89).

#### Effect of speech clarity on BOLD activity

Separately for each region of the Schaefer atlas, we analyzed the effect of speech masking on the BOLD activity to investigate whether story materials lead to similar neural activity changes compared to the activity changes reported previously for degraded/masked spoken sentences (Scott and Johnsrude, 2003; Scott et al., 2006; Wild et al., 2012a; Wild et al., 2012b; Scott and McGettigan, 2013; Ritz et al., 2022). For this analysis, a design matrix with 19 regressors was created. The design matrix contained one unique regressor for each of the speech-clarity conditions (clear speech, +14 dB, +9 dB, +4 dB, -1 dB SNR) for each of the three stories, one intercept regressor for each story, and one overall intercept regressor. The 15 speech-condition regressors were convolved with a canonical hemodynamic response function (Lindquist et al., 2009). One general linear model was calculated (Friston et al., 1994) for each participant and region of the Schaefer atlas, using the BOLD activity time course (concatenated for the three stories) as the predicted variable and the design matrix as the predictor. We examined the linear relationship between BOLD activity and speech-clarity conditions by multiplying a contrast vector with the estimated coefficients. Values of the contrast vector were coded as -2, -1, 0, 1, 2 for the clear speech, +14 dB, +9 dB, +4 dB, and -1 dB SNR conditions, respectively, whereas the other regressors were set to 0. This resulted in one contrast coefficient for each participant and region. Positive values indicate an increase in BOLD activity with increasing speech masking and associated listening effort, whereas negative values indicate an increase in BOLD signal with decreasing speech masking and associated intelligibility. For each Schaefer region, contrast coefficients were tested against zero using a one-sample t-test (group-level analysis) and the resulting t-value was converted to a z-score, mapped onto a partially inflated standard brain surface, and threshold at 3.89 (Bonferroni-corrected significance threshold).

#### Inter-subject correlation analysis

One important neural signature of the processing of naturalistic, continuous stimuli is the degree to which neural activity patterns are similar across participants, referred to as inter-subject correlation (ISC; Hasson et al., 2004; Hasson et al., 2010; Nastase et al., 2019; Regev et al., 2019). We focused on two types of ISC analyses, capitalizing on spatial and temporal pattern similarity (Nastase et al., 2019; Regev et al., 2019; Lee and Chen, 2022). Spatial ISC reveals the degree to which the neural activation patterns of different voxels within a brain region are shared among participants. Temporal ISC reveals the degree to which neural activation of a brain region evolves similarly over time across different individuals. ISC analyses were separately calculated for the three sub-groups of participants – that is, people who listened to the same randomization of speech-clarity conditions – before conducting group analyses involving all participants. For both spatial and temporal ISC, the whole BOLD signal time courses were first time-shifted by 5 s to account for the hemodynamic response delay (Buckner, 1998; Lindquist et al., 2009; Taylor et al., 2018).

For the spatial ISC analysis, the BOLD signal for each 30–33 s speech-clarity segment was averaged over time, separately for each voxel, and subsequently averaged across the segments with the same speech-clarity level in a story. The mean signal across the voxels within a Schaefer region was subtracted from the activity value of each voxel within that region (i.e., mean-centered), separately for each speech-clarity condition and story. The activity values for the three stories were subsequently concatenated, leading to one activity vector per Schaefer region, speech-clarity level, and participant. For each Schaefer region and speech-clarity level, a leave-one out procedure was implemented to calculate an ISC value for each participant. The activity vector of one participant was left out, and the activity vectors for the other n-1 participants were averaged. The activity vector of the participant who was left out was correlated with the averaged activity vector of the n-1 participants using Spearman correlation, and the resulting correlation value was used as the ISC value for the participant who was left out. The procedure was repeated such that each participant was left out once. The leave-one-out procedure was calculated separately for each sub-group of participants. These calculations resulted in one ISC value for each participant, speech-clarity condition, and brain region of the Schaefer atlas.

For the temporal ISC analysis, BOLD signal time courses for each Schaefer region were separated into individual 30–33 s speech-clarity segments. The mean BOLD signal for a given 30–33 s segment was subtracted from the BOLD signal at each sample of the segment (i.e., mean-centered). The mean-centered BOLD signal time courses of individuals segments and the 3 stories were concatenated, separately for each of the 5 speech-clarity conditions. For each Schaefer region, a leave-one out procedure was implemented to calculate an ISC value for each participant and speech-clarity condition. The concatenated time course of one participant was left out, and the time courses for the other n-1 participants were averaged. The time course of the participant who was left out was correlated with the averaged time course of the n-1 participants using Spearman correlation, and the resulting correlation value was used as the ISC value for the participant who was left out. The procedure was repeated such that each participant was left out once. The leave-one-out procedure was calculated separately for each sub-group of participants. These calculations resulted in one ISC value for each participant, speech-clarity condition, and Schaefer brain region.

To assess whether speech clarity affected spatial and temporal ISC, a linear function and quadratic fit was used to relate ISC values to speech-clarity conditions (coding -2, -1, 0, 1, 2 for the clear, +14 dB, +9 dB, +4 dB, and -1 dB SNR conditions, respectively). A linear and quadratic function was fit separately for each participant and brain region. A positive linear coefficient indicates higher ISC as speech masking increases (SNR decreases), whereas a negative linear coefficient indicates lower ISC values as speech masking increases. A positive quadratic coefficient indicates higher ISC for the two end points or one end point of the five speech-clarity conditions relative to the other conditions. The linear and quadratic coefficients were tested against zero using a one-sample t-test, separately for each brain region. The resulting t-value was converted to a z-score, mapped onto a partially inflated standard brain surface, and thresholded at 3.89 (Bonferroni-corrected significance threshold).

In some previous work using ISC, permutation analyses to calculate chance level ISC have been conducted (Honey et al., 2012). In the current study, the calculation of ISC z-scores for each participant and condition (instead of ISC values) by permuting spatial activation patterns and time-shifting temporal activation patterns, and subsequent statistical analyses for the ISC z-scores, led to qualitatively similar results as for the analyses described in the previous paragraphs. For simplicity and better interpretability of the results, we limit reporting of the results to analyses for ISC values and do not report ISC z-scores.

#### Activation to event boundaries

The times at which event boundaries occurred in a story were determined to investigate neural activation related to event boundaries. To this end, a transcription was obtained manually for each story. The transcription was used with OpenAI’s large language model GPT-4 (Generative Pre-trained Transformer 4; OpenAI et al., 2023) to identify event boundaries for each story. Previous work has shown that segmentation of stories into distinct events using OpenAI’s GPT closely aligns with event segmentation by humans (Michelmann et al., 2025; Panela et al., 2025). The segmentation approach was implemented in Python 3.11.5 (van Rossum and Drake, 2010) using OpenAI’s Application Programming Interface (API). The following prompt was input to OpenAI’s model to identify event boundaries (see also Michelmann et al., 2025; Panela et al., 2025): “An event is an ongoing coherent situation. The following story needs to be copied and segmented into large events. Copy the following story word-for-word and start a new line whenever one event ends and another begins. This is the story: …”. After this prompt, the full transcription of a story was inserted (without paragraph breaks or other formatting that could bias segmentation), followed by an additional prompt to refresh and reiterate the instructions: “This is a word-for-word copy of the same story that is segmented into large event units: ” (Michelmann et al., 2025; Panela et al., 2025). The temperature parameter of the model was set to 0 to facilitate a deterministic and reproducible output (Panela et al., 2025; max_tokens was set to 4096). Through this procedure, 44 events and corresponding boundaries were identified across the transcriptions of the three stories.

To obtain the times at which event boundaries occurred in the auditory story, we used the online implementation of Clarin’s forced alignment software. The forced alignment software provides the onset time for each word in a story using the story audio files and story transcriptions (https://clarin.phonetik.uni-muenchen.de/BASWebServices/; Kisler et al., 2017; incorrect estimations were manually corrected). The onset times of the words that OpenAI’s GPT-4 had identified as event boundaries (i.e., the beginning of a new event) were used for time-locking the BOLD signals for analysis. Two non-boundary control conditions were also created: “event center” and “sentence onset”. Times of the event-centers were calculated as the midpoint between two event boundaries. For the sentence-onset condition, we identified all non-boundary sentence-onset times from the output of Clarin’s forced alignment software (event boundaries typically coincide with sentence onsets). We reasoned that if neural responses to event boundaries reflect mental segmentation into meaningful story units, the responses should be larger than activity to event centers or non-boundary sentence onsets (Whitney et al., 2009).

Previous work shows a transient response peaking about 5 s following an event boundary (Speer et al., 2007; Zacks et al., 2010; Reagh et al., 2020), consistent with the current study. To capture the full response time course around event boundaries and the two control conditions, we identified for each Schaefer region the 20-second epochs in the BOLD time courses centered on the times of event boundaries, event centers, and non-boundary sentence onsets. Epochs started 10 s before and lasted for 10 s after and all epochs were averaged separately for event boundary, event center, or non-boundary sentence onset conditions; ignoring speech clarity for this analysis. The magnitude of the event-boundary response has previously been quantified by contrasting post- to pre-event boundary activity (Whitney et al., 2009). Following this approach, the BOLD signal across the -7 s to -1 s time window was averaged and served as a baseline (referred to as “pre” period) against which to contrast the averaged BOLD signal across the +1 s to +7 s time window (“post” period). Contrasts were created for the average post-minus-pre response for event boundary, event center, and non-boundary sentence onset conditions for each participant and brain region of the Schaefer atlas. For each Schaefer region, two paired samples t-tests were calculated, one testing whether the post-minus-pre response difference differed between event boundaries and event centers and the other testing whether the post-minus-pre response difference differed between event boundaries and non-boundary sentence onsets. The resulting t-values were converted to z-scores, mapped onto a partially inflated standard brain surface, and thresholded at 3.89 (Bonferroni-corrected significance threshold). Positive values indicate a larger response around event boundaries than event centers or non-boundary sentence onsets, whereas negative values indicate a smaller response.

To visualize time-locked response time courses for the different conditions (event boundary, event center, non-boundary sentence onset), the full (-10 to +10 s) response time courses were averaged across regions, separately for the 7 networks of the Schaefer atlas (Figure 1B; Yeo et al., 2011; Schaefer et al., 2018). For each network, we also examined differences in the time courses between responses to event boundaries and control conditions. Two paired-samples t-tests were calculated for each time point, contrasting event boundary vs event center and event boundary vs non-boundary sentence onset conditions. False discovery rate (FDR) was used to correct for multiple comparisons across time points (Benjamini and Hochberg, 1995; Genovese et al., 2002).

To investigate whether neural responses to event boundaries are affected by the level of speech masking, epochs were split into two groups depending on whether they occurred during high (clear, +14 dB SNR) or low speech-clarity segments (+4 dB, -1 dB SNR). Splitting speech-clarity conditions into two groups ensured that a sufficient number of event-boundary epochs per high (N=17, 16, 14 for the three versions of speech-clarity randomizations, respectively) and low (N=19, 21, 21) speech-clarity group were available for analysis. The BOLD signal across the -7 s to -1 s time window was averaged and subtracted from the averaged BOLD signal across the +1 s to +7 s time window, separately for high and low speech-clarity conditions. Then, the post-minus-pre activity difference for the low speech-clarity condition was subtracted from the post-minus-pre activity difference for the high speech-clarity condition, resulting in an interaction measure that reflects the change in event-boundary activation (post vs pre) with speech clarity. A positive value indicates a larger event-boundary response for high compared to low speech clarity. Grouping into low and high speech-clarity groups and similar calculation were also conducted for event centers (high: N=19, 20, 21; low: N=19, 16, 15; for the three versions of speech-clarity randomizations, respectively) and non-boundary sentence onsets (high: N=153, 168, 171; low: N=178, 146, 165). For statistical analyses, we compared the speech-clarity related change in event-boundary response with the speech-clarity related changes in event-center and sentence-onset responses using dependent samples t-tests, separately for each region of the Schaefer atlas. This contrast reflects the interaction between time window (post, pre), speech clarity (high, low), and condition (event boundary, event center, non-boundary sentence onset). T-values were converted to z-scores, mapped onto a partially inflated standard brain surface, and thresholded at 3.89 (Bonferroni-corrected significance threshold).

For each network of the Schaefer atlas, response time courses for the speech-clarity levels (high, low) and conditions (event boundary, event center, non-boundary sentence onset) were averaged across a network’s regions and used for visualization. BOLD signal differences between high and low speech-clarity levels were also examined for each network. A paired-samples t-test was calculated for each time point, contrasting high vs low speech-clarity levels, separately for event boundary, event center, and non-boundary sentence onset conditions. FDR was used to correct for multiple comparisons across time points (Benjamini and Hochberg, 1995; Genovese et al., 2002).

## Results

### BOLD signal changes with the changes in the speech envelope

To first confirm that stories in our paradigm can elicit meaningful activity, we examined how the BOLD signal responds to changes in the amplitude envelope of the clear speech signal. BOLD activity fluctuations in the superior temporal cortex were positively correlated with the fluctuations in the speech envelope (i.e., activity increased as the amplitude of the speech envelope increased; Figure 2A). In contrast, activity fluctuations in the left anterior insular cortex and bilaterally in the anterior cingulate cortex showed an anti-correlation with the speech-envelope fluctuations (i.e., activity increased with decreasing amplitude of the speech envelope; Figure 2A). The cingulo-opercular network is thought to subserve aspects of cognitive control and has been demonstrated to be sensitive to effort during listening (Herrmann et al., 2014; Henry et al., 2015; Eckert et al., 2016; Johnsrude and Rodd, 2016). The results may thus reflect increased effort associated with the comprehension of speech in babble when the speech signal is less intense. The speech-envelope analysis thus demonstrates that our story-listening paradigm can elicit meaningful changes in BOLD activity (cf. Whitney et al., 2009; Honey et al., 2012; Rowland et al., 2018).

**Figure 2:**
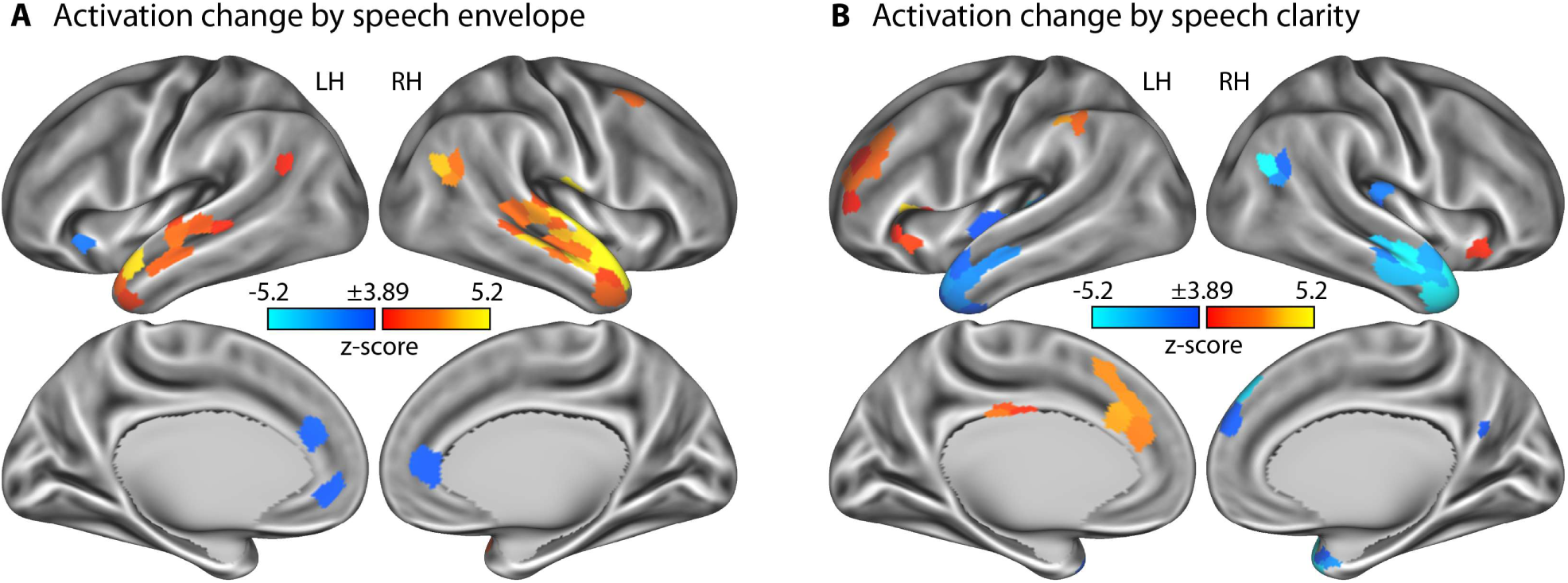
Neural activation associated with changes in the acoustic envelope of speech and speech-clarity conditions. **A:** Statistical z-score map for changes in BOLD signal as a function of the speech envelope. Positive values indicate a positive correlation, whereas negative values an anti-correlation between fluctuations in the speech envelope and the fluctuations in BOLD signal. **B:** Statistical z-score map for changes in BOLD signal as a function of speech-clarity conditions. Positive values indicate an increase in BOLD signal with decreasing speech clarity and associated listening effort, whereas negative values indicate an increase in BOLD signal with increasing speech clarity and associated intelligibility. Z-score maps are thresholded at a Bonferroni-corrected 0.05 significance level, corresponding to a z-score of 3.89. LH - left hemisphere; RH – right hemisphere.

### BOLD activity is sensitive to the clarity of speech during story listening

Next, we investigated whether our story-listening paradigm enables us to observe intelligibility- and effort-related activations that are commonly observed in studies using sentence materials (Davis and Johnsrude, 2003; Scott et al., 2006; Wild et al., 2012a; Wild et al., 2012b; Holmes and Johnsrude, 2021; Ritz et al., 2022). BOLD activity increased with decreasing speech masking (i.e., increasing intelligibility) bilaterally in auditory cortices and anterior temporal cortex, and in the right posterior temporal cortex (Figure 2B). In contrast, BOLD activity increased with increasing speech masking (i.e., potentially increasing effort) bilaterally in the anterior insula, the right dorsolateral prefrontal cortex, right supramarginal gyrus, and the right mid and anterior cingulate cortex (Figure 2B). The regions observed in these two contrasts are consistent with those identified in previous work using sentence materials, suggesting that BOLD activity in our story-listening paradigm is sufficiently sensitive to reveal intelligibility- and effort-related activations.

### Effect of speech clarity on spatial inter-subject correlation (ISC)

Spatial ISC reflects the degree to which the spatial activity patterns within a brain region are shared among listeners. Figure 3A shows that spatial ISC is relatively low for most speech-clarity conditions, considered on their own, with the exception of the most difficult, the -1 dB SNR, condition. For the -1 dB SNR condition, spatial ISC was strongest in the anterior insula, cingulate cortex, precuneus, and a few small regions in the temporal cortex.

**Figure 3:**
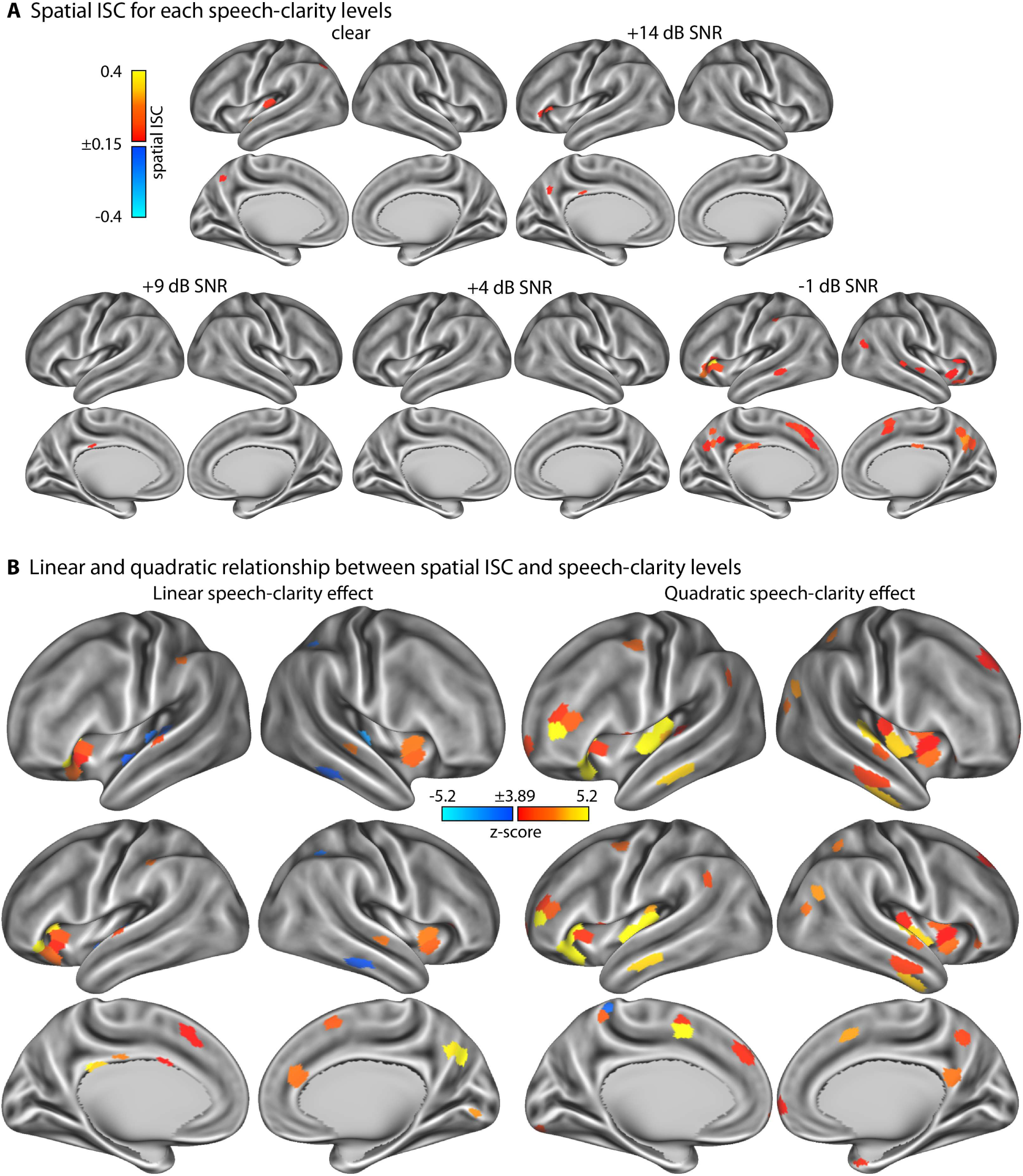
Results for spatial inter-subject correlation (ISC). **A:** Spatial ISC values for the five different speech-clarity conditions. **B:** Statistical z-score maps that reflect the linear (left) and quadratic (right) change in ISC as a function of speech-clarity condition. For the linear contrast, positive values indicate an increase in spatial ISC with decreasing speech clarity, whereas negative values indicate a decrease in spatial ISC with decreasing speech clarity. For the quadratic contrast, positive values indicate an increase in spatial ISC at either both ends or one end of the speech-clarity conditions relative to the moderately masked speech conditions. Z-score maps are thresholded at a Bonferroni-corrected 0.05 significance level, corresponding to a z-score of 3.89.

The linear function fit revealed an increase in spatial ISC – that is, spatial patterns of BOLD activity were more synchronized across listeners – in the precuneus, cingulate cortex, and anterior insula as speech clarity decreased (Figure 3B, left). The latter three regions are associated with listening effort (Wild et al., 2012b; Eckert et al., 2016; Johnsrude and Rodd, 2016; Peelle, 2018; Ritz et al., 2022). Spatial ISC decreased with decreasing speech clarity in a few auditory regions on the superior temporal plane and the temporal cortex (Figure 3B, left).

The analysis of quadratic spatial ISC trends also revealed a positive trend in the anterior insula, capturing the fact that spatial ISC increased in the anterior insula mainly in the most difficult condition (-1 dB SNR; Figure 3B, right). Spatial ISC also showed a quadratic trend in auditory regions, capturing a specific increase in spatial ISC for clear speech (compare Figure 3A and Figure 3B, right). These changes in spatial ISC were independent of overall activity differences in these regions, because the mean BOLD activity was removed prior to ISC analyses (see Methods).

### Effect of speech clarity on temporal inter-subject correlation (ISC)

Temporal ISC reflects the degree to which the BOLD activity time courses evolve similarly – that is, are synchronized – across listeners. Figure 4A shows that activity is most strongly synchronized across listeners in bilateral superior temporal cortices and inferior frontal cortex for all speech-clarity conditions, and in the precuneus in all but the most difficult condition (-1 dB SNR).

**Figure 4:**
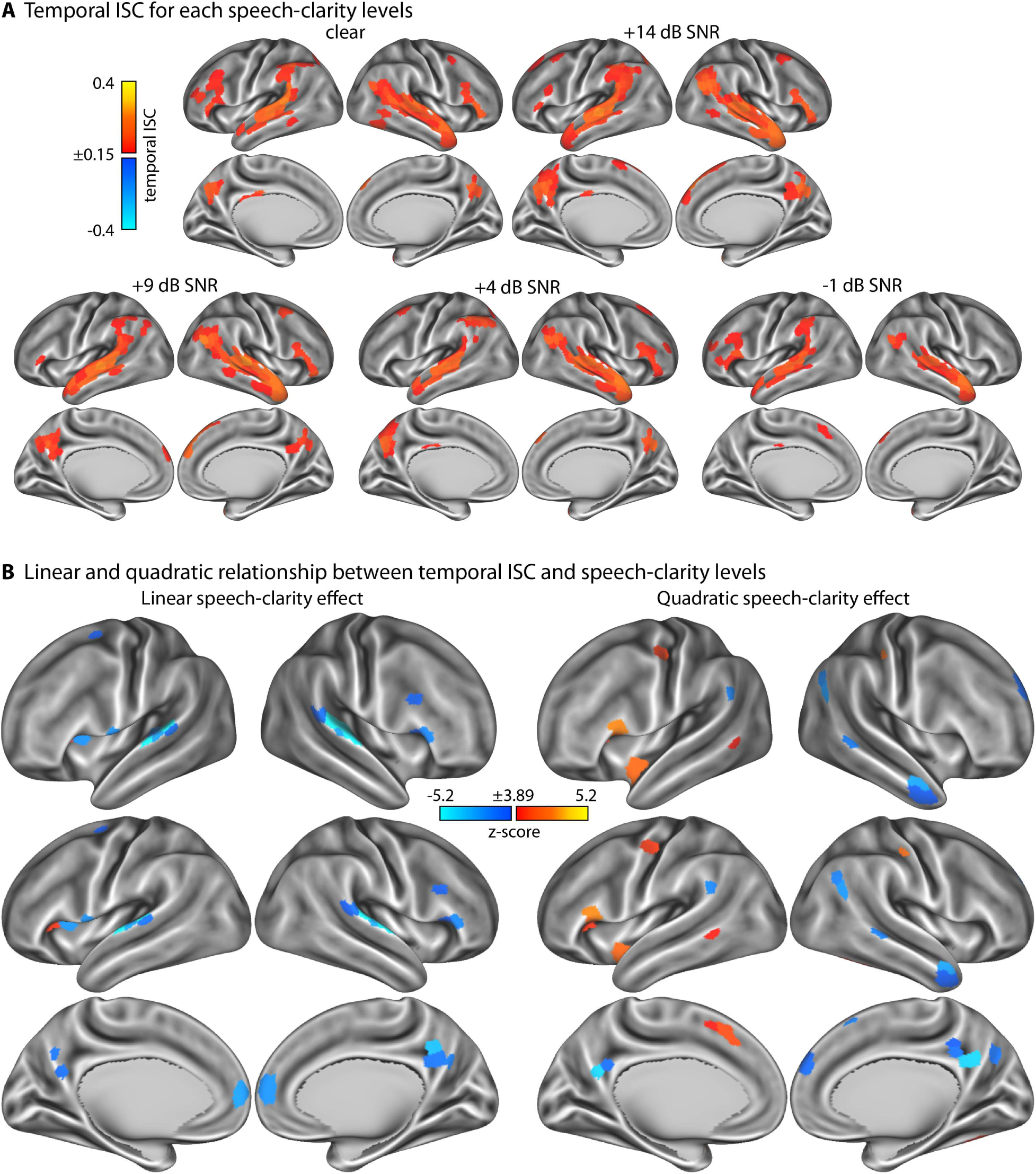
Results for temporal inter-subject correlation (ISC). **A:** Temporal ISC values for the five different speech-clarity conditions. **B:** Statistical z-score maps that reflect the linear (left) and quadratic (right) change in ISC as a function of speech-clarity condition. For the linear contrast, positive values indicate an increase in temporal ISC with decreasing speech clarity, whereas negative values indicate a decrease in temporal ISC with decreasing speech clarity. For the quadratic contrast, positive values indicate an increase in temporal ISC at either both ends or one end of the speech-clarity conditions relative to the moderately masked speech conditions. Z-score maps are thresholded at a Bonferroni-corrected 0.05 significance level, corresponding to a z-score of 3.89.

The linear function fit revealed a decrease in temporal ISC with decreasing speech clarity bilaterally in the superior temporal plane, precuneus, dorsal anterior cingulate and anterior prefrontal cortex, and the inferior frontal opercular cortex, including parts of the anterior insula (Figure 4B, left). There was also a region in the left anterior insula that showed an increase in temporal ISC with decreasing speech masking (Figure 4B, left), but ISC values in this region were overall very low (Figure 4A).

The quadratic trend analysis revealed a positive trend for temporal ISC in relation to speech-clarity levels in the left dorsal cingulate cortex extending to the supplementary motor area, consistent with the high temporal ISC in these regions specifically for the most difficult SNR (-1 dB; compare Figure 4A and Figure 4B, right). Additional positive quadratic trends were observed in the left inferior frontal cortex that seemed to be driven by the higher temporal ISC for the clear condition and the most difficult SNR compared to intermediate speech-clarity levels. Temporal ISC values were very low for the other few regions showing a quadratic trend (compare Figure 4A and Figure 4B, right). The right precuneus and a few regions along the temporal cortex showed a negative quadratic trend, driven mainly by the reduced temporal ISC for the most difficult speech-clarity condition (particularly the precuneus). These changes in the temporal synchronization of neural activation across listeners are independent of overall activation differences, because the mean BOLD signal was subtracted prior to ISC analyses.

### Transient responses around event boundaries during story listening

Neural responses to event boundaries reflect an important neural signature of event segmentation during the encoding of natural, continuous environments (Speer et al., 2007; Whitney et al., 2009; Zacks et al., 2010). We investigated changes in BOLD activity associated with event boundaries during continuous speech relative to two control conditions: event centers and non-boundary sentence onsets. This analysis was conducted first across all speech-clarity conditions to establish which brain regions respond to event boundaries. Figures 5A and B show the statistical z-score maps for differences in neural activation between event boundaries and event centers or non-boundary sentence onsets, respectively.

**Figure 5:**
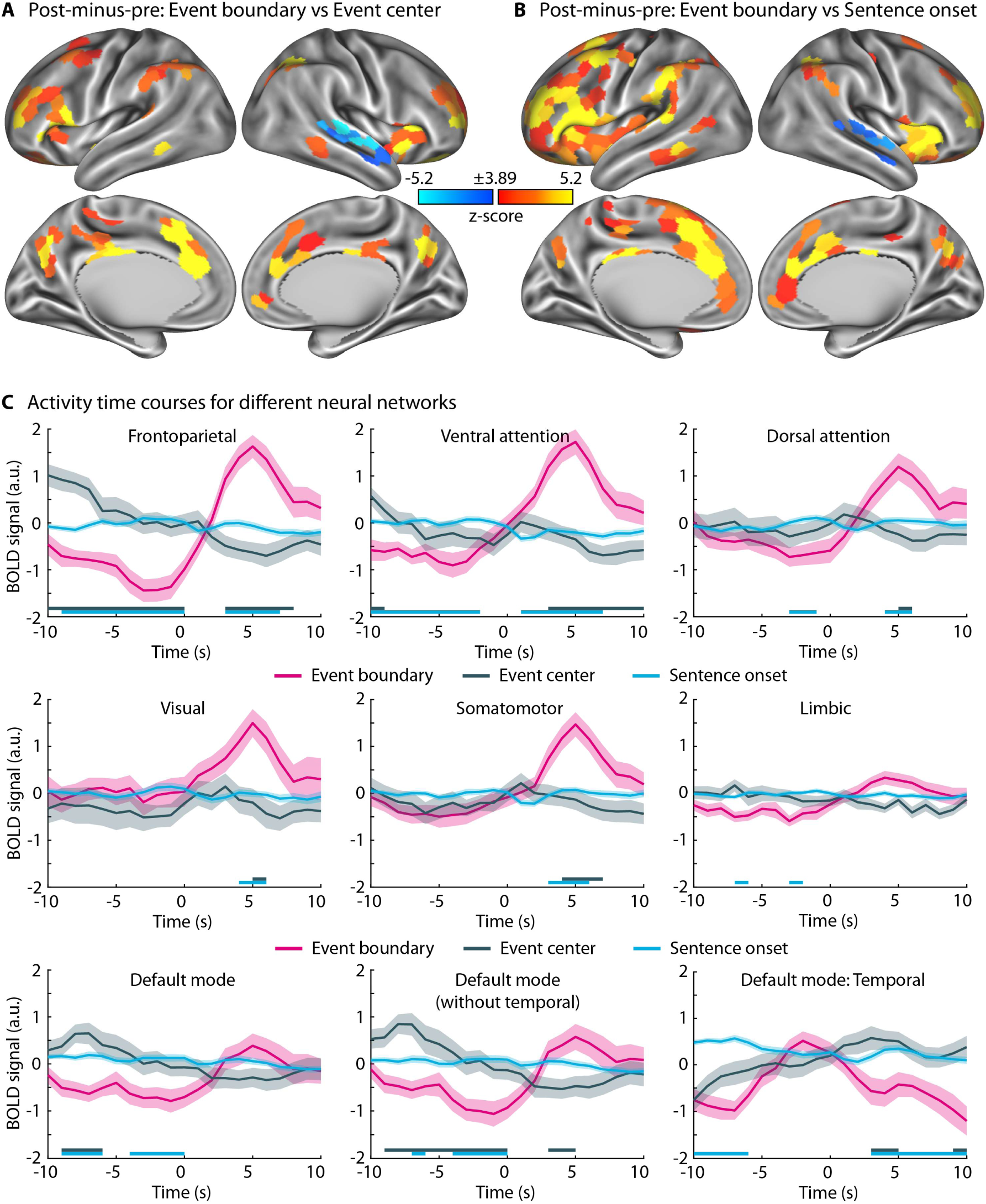
Neural activity at event boundaries. **A and B:** Statistical z-score maps showing the differences in neural activation between event boundaries and event centers (panel A) or non-boundary sentence onsets (panel B). The difference between the post- and the pre-time-locked windows (+1 to +7 s minus -7 to -1 s) was used as the neural activation signal for the displayed analyses. Positive values reflect a larger activation for event boundaries, whereas negative values reflect a smaller activation for event boundaries than for event centers or non-boundary sentence onsets. Z-score maps are thresholded at a Bonferroni-corrected 0.05 significance level, corresponding to a z-score of 3.89. **C:** BOLD signal time courses for 7 different networks of the Schaefer brain atlas time-locked to event boundaries, event centers, or non-boundary sentence onsets. Activation time courses for the default-mode network, excluding the temporal region, and for only the temporal region of the network to better display the negative effects in temporal cortex. The shaded areas around the mean BOLD time course reflect the standard error of the mean. Solid lines close to the x-axis indicate a significant difference between event boundary vs event center or event boundary vs non-boundary sentence onset (FDR-thresholded).

Neural activations were larger for event boundaries compared to event centers and non-boundary sentence onsets in several brain regions that are part of the frontoparietal, ventral and dorsal attention, visual, somatomotor, and parts of the default-mode networks (Figure 5C). The frontoparietal and ventral attention networks appear to be particularly responsive to event boundaries. Activation increased and peaked at around 5 s following an event boundary. The temporal evolution of the activation in the frontoparietal network also included an activation decrease prior to the event boundary (Figure 5C, top left).

In addition to the large number of regions showing greater neural activation for event boundaries than control conditions, the right temporal cortex exhibited smaller neural activation for event boundaries than event centers and non-boundary sentence onsets (Figures 5A and 5B). This effect appears to be driven by an activation peak just prior to an event boundary that was absent for event centers and non-boundary sentence onsets (Figure 5C, bottom right), resulting in the negative activation defined as the difference between the post- and the pre-time locked windows: +1 to +7 s minus -7 to -1 s.

### Change in event-boundary response associated with speech clarity

To investigate how event-boundary activations are affected by speech clarity, we grouped the speech-clarity conditions according to high and low clarity. Neural activation was investigated as the difference between the BOLD signal in the post- and the pre-time-locked windows: +1 to +7 s minus - 7 to -1 s. Figure 6 shows neural activation to event boundaries relative to event centers or non-boundary sentence onsets, separately for high and low speech clarity. Event boundaries elicited neural activation in the anterior insula, inferior frontal cortex, anterior cingulate cortex, dorsolateral pre-frontal cortex, precuneus, posterior temporal cortex, and parietal cortex for both high and low speech clarity (Figure 6, top row), although fewer regions were significant for low than high speech clarity (Figure 6, middle and bottom row, respectively).

**Figure 6:**
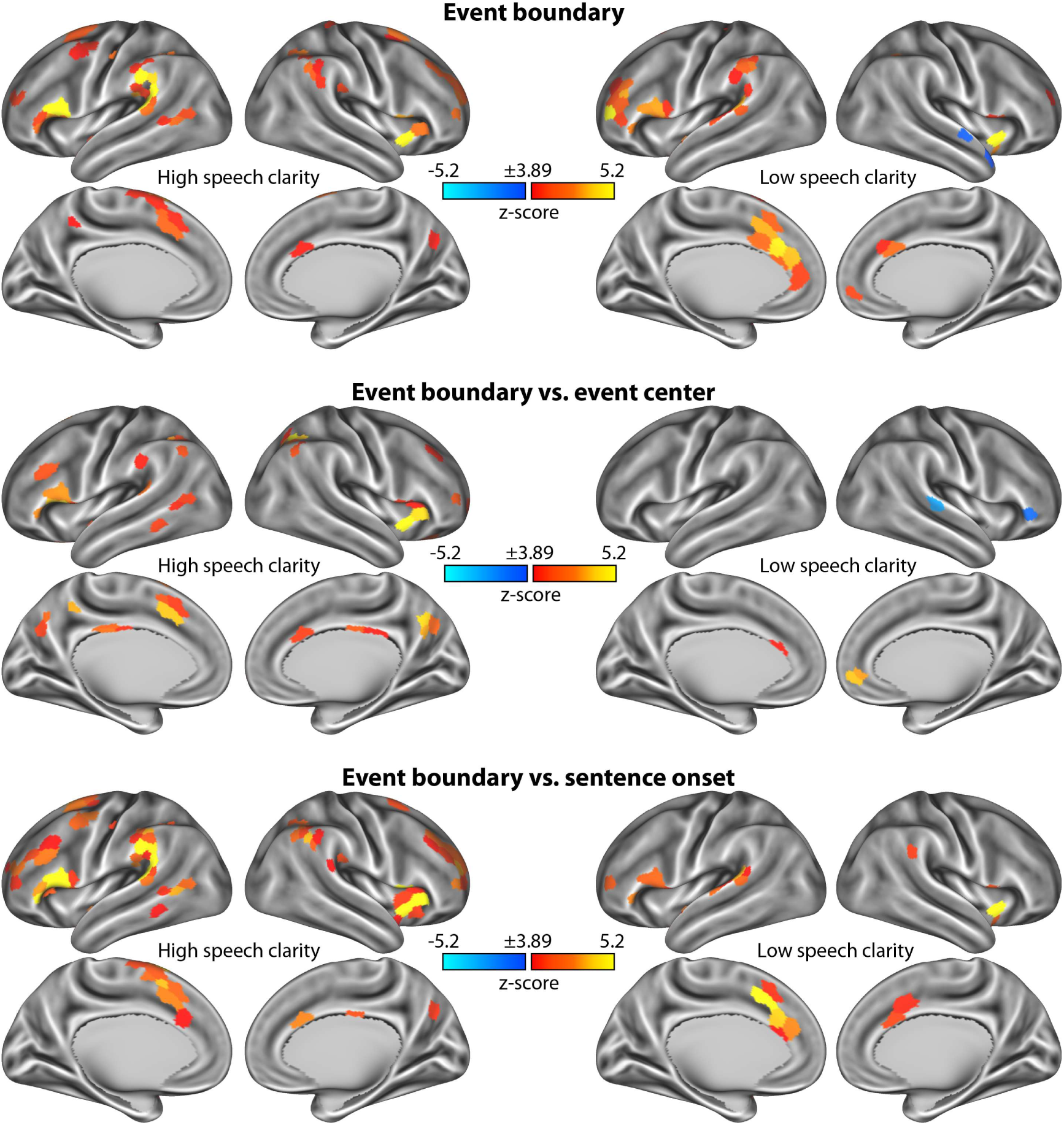
Neural activity at event boundaries, separately for low and high speech clarity. **Top row:** Statistical z-score maps showing the neural activation associated with event boundaries for high and low speech clarity. Neural activation was defined as the difference between the post- and the pre-event-boundary window (+1 to +7 s minus -7 to -1 s). Z-score maps resulted from testing the neural activation against zero. **Middle row:** Statistical z-score maps showing the differences in neural activation (post-minus-pre difference) between event boundaries and event centers. Positive values reflect a larger activation for event boundaries, whereas negative values reflect a smaller activation for event boundaries than for event centers. **Bottom row:** Same as in the middle row, contrasting activations to event boundaries relative to non-boundary sentence onsets. Z-score maps are thresholded at a Bonferroni-corrected 0.05 significance level, corresponding to a z-score of 3.89.

Directly contrasting responses for high compared to low speech clarity did not reveal a significant difference in any of the regions of the Schaefer atlas. The time courses for each brain network displayed in Figure 7 indicate, in line with Figure 6, that event boundaries elicit activation for both low and high speech-clarity conditions, suggesting that listeners can identify meaningful units in speech even in the presence of background masking. Activity was greater for high compared to low SNRs around event centers in the default mode network and for non-boundary sentence onsets in the limbic network, which overlap with the overall speech-clarity effect shown in Figure 2.

**Figure 7:**
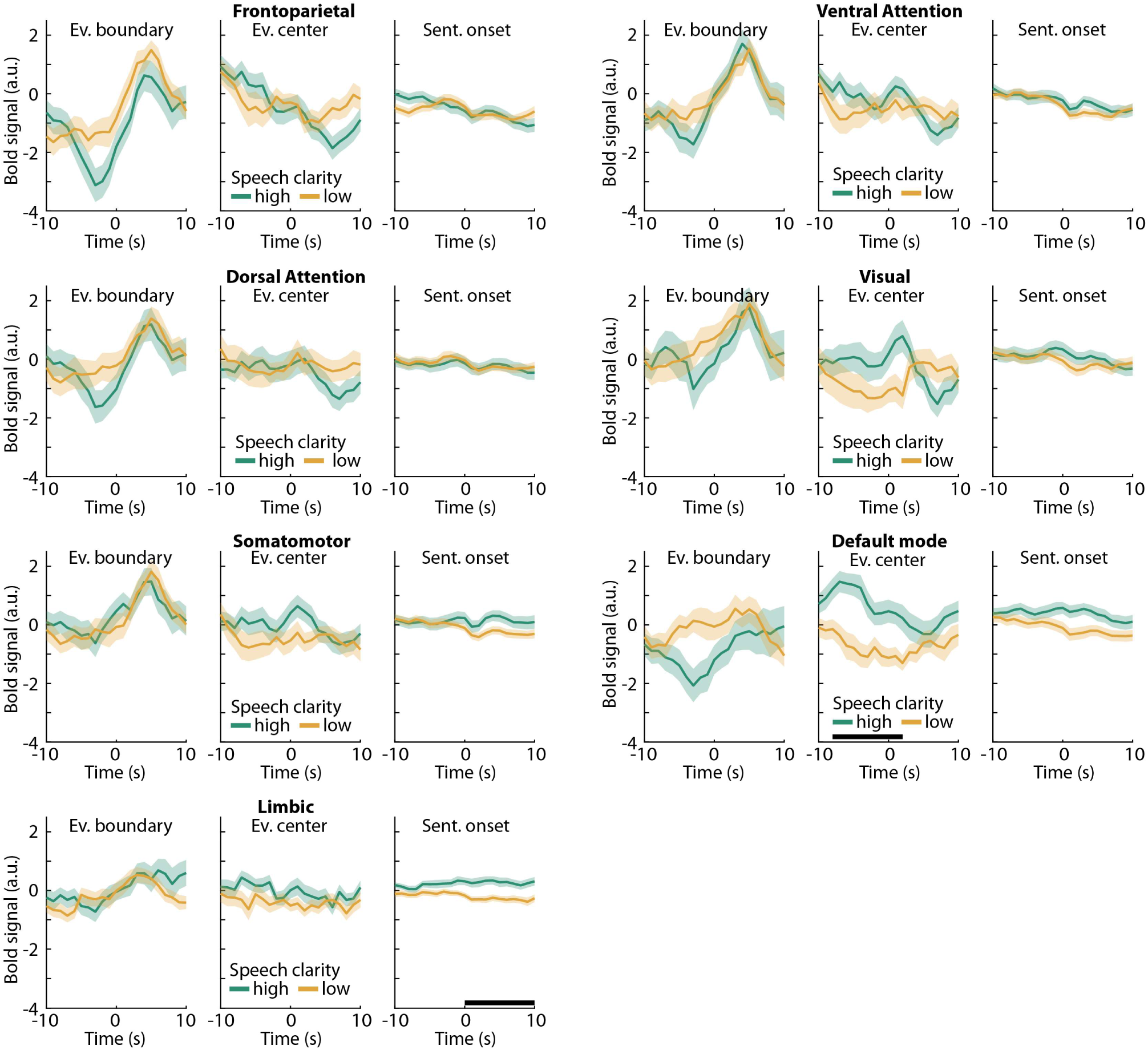
Network time courses for event boundary responses for high versus low speech clarity. BOLD signal time courses for 7 different networks of the Schaefer brain atlas. Separate time courses for high and low speech-clarity levels are shown. The shaded areas around the mean BOLD time course reflect the standard error of the mean. A black solid line close to the x-axis indicates a period during which high vs low speech-clarity conditions differ significantly (FDR-thresholded).

## Discussion

The current fMRI study investigated how neural signatures of continuous story listening change in the presence of multi-talker background babble. We show that neural activation increases in the anterior insula, anterior cingulate cortex, dorsolateral prefrontal cortex, and supramarginal gyrus when speech masking increases (Figure 2). Temporal activation patterns in auditory brain regions became more idiosyncratic, that is, less similar, across listeners with increasing speech masking (Figure 4). In contrast, spatial (and to some extent temporal) activation patterns were more similar across listeners in the anterior insula, anterior cingulate, precuneus, and dorsolateral prefrontal cortex when participants listened to minimally intelligible compared to highly intelligible speech, indicating shared neuro-cognitive processes (Figure 3). We further observed increased activation most prominently in the frontoparietal, ventral, and dorsal attention networks at times during story listening when one meaningful event ended and another began (Figure 5). This activation to event boundaries appeared to be little affected by background masking (Figures 6 and 7). In sum, neural signatures of story listening indicate more idiosyncratic processing of the acoustic information in auditory regions, but shared neural processes and intact gist processing when stories are masked by background babble.

### Neural activation associated with changes in the speech envelope and background masking

Neural activation was modulated by the fluctuations in the amplitude envelope of the speech signal, such that activity in the superior temporal cortex, putative auditory cortex near Heschl’s gyrus, increased with increasing speech amplitude (Figure 2A). Auditory activity correlated with changes in the speech envelope have also been reported previously in studies using fMRI (Honey et al., 2012), functional near-infrared spectroscopy (Rowland et al., 2018), and electro-/magnetoencephalography (Lalor and Foxe, 2010; Ding and Simon, 2013; Ding et al., 2014; Vanthornhout et al., 2018; Panela et al., 2024), likely reflecting the acoustic processing of the speech.

We further observed that activity increased in the anterior insula and anterior cingulate cortex with decreasing amplitude of the speech signal (i.e., showing anti-correlated fluctuations between neural activity and the speech envelope; Figure 2A). The anterior insula and cingulate cortex, in addition to the dorsolateral prefrontal cortex, were also activated when the speech was masked by background babble (Figure 2B). The cingulo-opercular network, including the insula, cingulate cortex, prefrontal cortex, and parietal cortex are frequently observed when individuals engage in challenging tasks, including listening to degraded speech and identifying subtle changes in non-speech sounds (Dosenbach et al., 2006; Dosenbach et al., 2008; Wild et al., 2012b; Scott and McGettigan, 2013; Herrmann et al., 2014; Henry et al., 2015; Eckert et al., 2016; Johnsrude and Rodd, 2016; Peelle, 2018; Ritz et al., 2022). Increased activity in the cingulo-opercular network is thought to reflect increased executive control demands during challenging tasks (Dosenbach et al., 2006; Cole and Schneider, 2007; Dosenbach et al., 2008; Crittenden et al., 2016; Gratton et al., 2018; Hausman et al., 2022; Dosenbach et al., 2025), and we extend this here to story listening, demonstrating that this effort response may fluctuate moment to moment, reflecting the dynamic demands of the listening situation.

We further found increased activation in the anterior temporal cortex, precuneus, and posterior temporal cortex as speech-clarity and hence intelligibility increased (Figure 2B), which is consistent with previous sentence-listening paradigms (Narain et al., 2003; Scott et al., 2006; Obleser et al., 2007; Obleser et al., 2008; Wild et al., 2012a; Evans et al., 2016; Holmes and Johnsrude, 2021) and with the idea that these areas subserve suggested language-specific processes (Fedorenko and Thompson-Schill, 2014; Fedorenko et al., 2024).

The current data thus show that a listening paradigm with ∼10 min stories can reveal activations that are comparable to more traditional sentence-listening paradigms (see Evans et al., 2016 for ∼20 s narratives), with the advantage that stories are more enjoyable for listeners and enable investigating questions about time-varying and more naturalistic speech processes.

### Changes in shared neural activity patterns as speech masking increases

The current study investigated the extent to which individuals share neural activity patterns while listening to spoken stories in the presence of background babble. We focused on the similarity of the temporal evolution of neural activity and the spatial similarity of neural activity profiles across voxels within a brain region (temporal and spatial inter-subject correlation, respectively; Hasson et al., 2010; Nummenmaa et al., 2018; Nastase et al., 2019).

Temporal ISC was greatest along the superior temporal cortex, inferior frontal cortex, and precuneus for all speech-clarity conditions, consistent with previous work on story listening under clear conditions (Honey et al., 2012; Regev et al., 2019). Temporal ISC decreased with increasing speech masking in auditory regions in the superior temporal plane (Figures 4). Reduced temporal ISC in the superior temporal plane may reflect more idiosyncratic processing and temporal tracking of the acoustic speech properties (Honey et al., 2012; Rowland et al., 2018). Temporal ISC also decreased in the posterior medial cortex (precuneus) and medial prefrontal cortex as masking level increased. Both of these regions have been linked to engagement and shared experiences with movies or spoken narratives (Lerner et al., 2011; Schmälzle et al., 2015; Simony et al., 2016; Nguyen et al., 2019; Song et al., 2021; Stawarczyk et al., 2021), potentially suggesting the experiential tracking of the narrative over time becomes more idiosyncratic when speech is highly masked, perhaps reflecting individual differences in compensation for degradation. Overall, however, temporal ISC was distributed widely over the cortex for all speech-clarity conditions, and this is consistent with attentive story listening (Regev et al., 2019) and story comprehension (Honey et al., 2012).

Spatial ISC was relatively low for all speech-clarity conditions considered individually, except for the most difficult SNR (Figure 3) and in auditory regions for clear speech. That only clear speech, but none of the babble-masked speech conditions, yielded reliable spatial ISC in auditory regions is consistent with the increased idiosyncratic activity pattern in auditory regions observed also for temporal ISC. Interestingly, spatial (and to some extent temporal) ISC increased in the precuneus, anterior insula, and anterior cingulate with increasing speech masking, particularly for the minimally intelligible condition (-1 dB SNR). The latter two brain regions comprise the “cingulo-opercular network” which has frequently been implicated in executive, cognitive control and with listening effort during speech comprehension (Figure 2B; Wild et al., 2012b; Herrmann et al., 2014; Henry et al., 2015; Eckert et al., 2016; Johnsrude and Rodd, 2016; Ritz et al., 2022). The increase in spatial ISC in the cinguloopercular regions may indicate a similar mode of processing challenging speech. ISC in the posterior medial cortex (precuneus) has been linked to engagement and shared experiences (Schmälzle et al., 2015; Simony et al., 2016; Nguyen et al., 2019; Song et al., 2021; Stawarczyk et al., 2021). The increase in spatial ISC in the precuneus is surprising given the concurrent decrease in temporal ISC in this region, although the two appear to spatially dissociate somewhat along the anterior (temporal ISC) to posterior (spatial ISC) axis. We speculate that the decrease in temporal ISC reflects more idiosyncratic narrative tracking over time, whereas the increase in spatial ISC may be due to similar tonic attentional engagement across participants, as the difficulty to comprehend speech increased.

### Neural responses to event boundaries are little affected by speech masking

We observed increased neural activation in a large number of parcellated brain regions, covering frontoparietal, ventral attention, and dorsal attention networks, for story event boundaries relative to event centers and non-boundary sentence onsets (Figure 5). This is consistent with previous research demonstrating event-boundary related increases in neural activation during movie watching, narrative reading, and story listening (Speer et al., 2007; Whitney et al., 2009; Zacks et al., 2010; Ben-Yakov and Henson, 2018; Reagh et al., 2020; Stawarczyk et al., 2021). The activation increase at event boundaries is thought to index the updating of mental representations at an event boundary (Speer et al., 2007; Whitney et al., 2009; Zacks et al., 2010) and the magnitude of the event-boundary response has been linked to event memory (Ben-Yakov and Dudai, 2011).

The temporal evolution of the activations around event boundaries were characterized by relatively lower activity prior to event boundaries (compared to the event centers and non-boundary sentence onsets), particularly in the frontoparietal network, and peak activity about 5 s after event boundary onsets. The pre-boundary activity decrease and the hemodynamic response delay of about 4-5 s (Buckner, 1998; Lindquist et al., 2009; Taylor et al., 2018) suggest that little or no information about the new event is available to participants by the time the neural response is elicited. This may suggest that participants use prior experiences with stories and information from event endings to predict event boundaries during story listening, and in turn drive the response, rather than relying on the information following the onset of a new event. Preliminary work from our lab indeed suggests that participants identify event boundaries when they recognize a meaningful unit ending rather than a new meaningful unit beginning during story listening (Lamekina et al., in prep).

Event boundaries are often identified through human raters (Zacks et al., 2006; Kurby and Zacks, 2011; Sargent et al., 2013; Kurby et al., 2014; Lee and Chen, 2022; Pitts et al., 2022; Sasmita and Swallow, 2022), but recent research suggests that modern large language models (LLMs) can identify event boundaries in stories similarly well (Michelmann et al., 2025; Panela et al., 2025). Our observation of larger activation to LLM-identified event boundaries than to event centers and non-boundary sentence onsets (Figure 5) provides additional evidence that event boundaries identified through large language models are meaningful to human listeners as indexed through their brain responses.

Although listening to the stories under low speech clarity led to fewer regions that significantly responded to event boundaries compared to listening under high speech clarity (Figure 6), we did not observe a significant difference when directly comparing the event boundary response between high and low speech clarity (Figure 7). Many brain regions in frontal cortex and in midline structures exhibited an event-boundary response for both high and low speech clarity (Figure 6), potentially suggesting that background masking does not overly affect story comprehension and thus event-boundary related processes. We have previously shown in behavioral and neural work that individuals remain engaged and find stories of the kind played here absorbing even when speech is presented in background noise (Herrmann and Johnsrude, 2020a; Irsik et al., 2022a), and intelligibility (the ability to report words from sentences) is reduced. The activation increase for event boundaries was present even for speech at low speech clarity levels (that lead to more than 20% reduced speech intelligibility; Irsik et al., 2022a), suggesting that individuals can identify meaningful units under such conditions, which perhaps helps them to stay engaged in listening when speech is interesting to them.

## Conclusions

The current fMRI study reveals a diverse set of changes in brain activation patterns while individuals listened to naturalistic, spoken stories masked by different degrees of multi-talker background babble noise. Spatial and temporal activation patterns in auditory regions became more idiosyncratic to individual listeners as background babble increased. However, spatial activation patterns were more similar across participants in brain regions associated with executive control and effort, indicating that shared processes are recruited during challenging listening. We further show that neural activation at the boundaries between meaningful events in a story increased in a large number of brain regions, most prominently in the frontoparietal and dorsal/ventral attention networks. There was little evidence of a decline in the event-boundary response when speech was moderately masked by background babble. Overall, the current results suggest that although the processing of acoustic information is more idiosyncratic under background babble, processing of a story’s meaning is little affected by background masking (as indicated by event-boundary response), even when listening is effortful. The data may indicate that people stay engaged during story listening and that several cognitive processes associated with speech processing – potentially including processes reflecting engagement and enjoyment related to anticipation, suspense, and empathy – remain intact even under background noise.

## Author Contributions

**BH:** Conceptualization, methodology, validation, formal analysis, data curation, writing – original draft, writing – review and editing, visualization, project administration, supervision. **AM:** Conceptualization, methodology, investigation, data curation, writing – review and editing, project administration. **RAP:** Formal analysis, writing – review and editing. **ISJ:** Conceptualization, methodology, writing – review and editing, project administration, supervision, funding acquisition.

## Data Availability

Consent for public data sharing was not obtained from participants when study data were recorded, and the data thus cannot be made publicly available.

## Statements and Declarations

The authors have no conflicts or competing interests.

## Acknowledgements

BH was supported by a BrainsCAN postdoctoral fellowship (Canada First Research Excellence Fund; CFREF) and by the Canada Research Chair program (CRC-2023-00383). AM was supported by a postdoctoral fellowship from the Canadian Institutes of Health Research (CIHR). RAP was supported by a doctoral scholarship from CIHR (193310). The research was supported by funding from the CIHR awarded to ISJ (470281) and BH (517611). Neuroimaging was supported by the Centre for Functional and Metabolic Mapping Internal Funding Program and a Canada First Research Excellence Foundation (CFREF) grant.

